# Immunogenic Potential of DNA Vaccine candidate, ZyCoV-D against SARS-CoV-2 in Animal Models

**DOI:** 10.1101/2021.01.26.428240

**Authors:** Ayan Dey, T.M. Chozhavel Rajanathan, Harish Chandra, Hari P.R. Pericherla, Sanjeev Kumar, Huzaifa S. Choonia, Mayank Bajpai, Arun K. Singh, Anuradha Sinha, Gurwinder Saini, Parth Dalal, Sarosh Vandriwala, Mohammed A. Raheem, Rupesh D. Divate, Neelam L. Navlani, Vibhuti Sharma, Aashini Parikh, Siva Prasath, Sankar Rao, Kapil Maithal

## Abstract

Severe Acute Respiratory Syndrome Coronavirus 2 (SARS-CoV-2), initially originated in China in year 2019 and spread rapidly across the globe within 5 months, causing over 96 million cases of infection and over 2 million deaths. Huge efforts were undertaken to bring the COVID-19 vaccines in clinical development, so that it can be made available at the earliest, if found to be efficacious in the trials. We developed a candidate vaccine ZyCoV-D comprising of a DNA plasmid vector carrying the gene encoding the spike protein (S) of the SARS-CoV-2 virus. The S protein of the virus includes the receptor binding domain (RBD), responsible for binding to the human angiotensin converting enzyme (ACE-2) receptor. The DNA plasmid construct was transformed into *E. coli* cells for large scale production. The immunogenicity potential of the plasmid DNA has been evaluated in mice, guinea pig, and rabbit models by intradermal route at 25, 100 and 500μg dose. Based on the animal studies proof-of-concept has been established and preclinical toxicology (PCT) studies were conducted in rat and rabbit model. Preliminary animal study demonstrates that the candidate DNA vaccine induces antibody response including neutralizing antibodies against SARS-CoV-2 and also provided Th-1 response as evidenced by elevated IFN-γ levels.

## Introduction

Three highly pathogenic human coronaviruses (CoVs) have been identified so far, including Middle East respiratory syndrome coronavirus (MERS-CoV), severe acute respiratory syndrome coronavirus (SARS-CoV) and severe acute respiratory syndrome coronavirus-2 (SARS-CoV-2). Among them, SARS-CoV was first reported in Guangdong, China in 2002 ^1^. SARS-CoV caused human-to-human transmission and resulted in the 2003 outbreak with about 10% case fatality rate (CFR), while MERS-CoV was reported in Saudi Arabia in June 2012 ^2^. Even though with its limited human-to human transmission, MERS-CoV showed a CFR of about 34.0 %^3^. The SARS-CoV-2 was first reported in Wuhan, China in December 2019 from patients with pneumonia, and it has exceeded both SARS-CoV and MERS-CoV in its rate of transmission among humans^4,5^. The outbreak of a novel coronavirus (SARS-CoV-2) represents a pandemic threat that has been declared a public health emergency of international concern (PHEIC). Currently, the intermediate host of SARS-CoV-2 is still unknown, and no effective prophylactics or therapeutics are available, though various drugs have shown mild to moderate protection but none of them have shown conclusive evidence.

Several pre-clinical or clinical trials are going on, which include repurposing of already approved drugs but with different indications such as anti-malarial, anti-viral, anti-parasitic drugs, or monoclonal antibodies, etc^6,7^. However, these drugs may help to prevent worsening of the coronavirus infection only and there is still an unmet need of a vaccine against novel coronavirus SARS-CoV-2.

Huge progresses were made in last one year for bringing effective vaccines against SARS-CoV-2. As per WHO draft landscape of vaccines, currently 172 vaccine candidates are in pre-clinical development, 63 are in clinical development. Among the candidates in clinical development, 6 are based on plasmid DNA technology including our vaccine candidate. Other 5 candidates are from Inovio Pharmaceuticals/Beijing Advaccine Biotechnology/VGXI Inc./ Richter-Helm BioLogics/Ology Bioservice; Osaka University/ AnGes/ Takara Bio/ Cytiva; Genexine Consortium (GenNBio, International Vaccine Institute, Korea Advanced Institute of Science and Technology (KAIST), Pohang University of Science and Technology (POSTECH)/ Binex/ PT Kalbe Pharma, Providence Health & Services. Further, 14 candidates of total 162 are based on DNA vaccine platform and are in preclinical development^8^.

Three candidates including two mRNA based candidate from Pfizer and Moderna and Chimpanzee adenovirus vector based candidate from AstraZeneca got emergency use approval globally. The emergency use was approved based on Phase-3 efficacy data. The mRNA vaccines from Pfizer reported 95% efficacy^9^, whereas Moderna and AstraZeneca reported 94.5 % and 70.4% efficacy respectively for their vaccine candidate^10^.

The conventional active vaccines are made of a killed or attenuated form of the infectious agent. Vaccination with live attenuated and killed vaccines in most cases results in generation of humoral but not a cell-mediated immune response. What is required in such cases, but not available, are antigens that are safe to use, that can be processed by the endogenous pathway and eventually activating both B and T cell response. The activated lymphocytes generated would destroy the pathogen-infected cell. For these reasons, a new approach of vaccination that involves the injection of a piece of DNA that contains the genes for the antigens of interest are under investigation. DNA vaccines are attractive because they ensure appropriate folding of the polypeptide, produce the antigen over long periods, and do not require adjuvants. These host-synthesized antigens then can become the subject of immune surveillance in the context of both major histocompatibility complex class I (MHC I) and MHC II proteins of the vaccinated individual^11^. By contrast, standard vaccine antigens are taken up into cells by phagocytosis or endocytosis and are processed through the MHC class II system, which primarily stimulates antibody response. In addition to these properties, the plasmid vector contains immunostimulatory nucleotide sequences-unmethylated cytidine phosphate guanosine (CpG) motifs - that induce strong cellular immunity^12^. Finally, DNA vaccines have been shown to persist and stimulate sustained immune responses. Other advantages are that the technology for producing the vaccine is very simple and rapid, secondly the DNA molecule is stable, has a long shelf life, and does not require a strict cold chain for distribution. DNA vaccines are also safer than certain live-virus vaccines, especially in immunocompromised patients. It also circumvents the numerous problems associated with other vaccines, such as immune responses against the delivery vector and concern about safety related to the use of any viral vector^13^.

Prior studies have demonstrated that a DNA vaccine approach for SARS and MERS can induce immune response including neutralizing antibody (nAb) responses in clinical trials and provide protection in challenge models. Previous studies indicated immunization in animal models with DNA vaccines encoding MERS-CoV spike (S) protein provided protection against disease challenge with the wild type virus. In subjects immunized with MERS-CoV DNA vaccine durable neutralizing antibodies (nAbs) and T cell immune responses were measured, and a seroconversion rate of 96% was observed and immunity was followed for 60 weeks in most study volunteers^14^. Similarly, NIH completed Phase 1 clinical trial for SARS-DNA vaccine. Dose of 4.0 mg was tested in healthy adults who were vaccinated on days 0, 28, and 56. The vaccine was found to be well-tolerated and induced antibody responses against the SARS-CoV in 80% of subjects after 3 doses^15^. More recently, Inovio pharmaceuticals developed DNA vaccine INO-4800 against SARS-CoV-2, which was found to be safe and immunogenic in Phase-I trial, eliciting either or both humoral or cellular immune responses^16^.

The spike proteins of SARS-CoV-2 and SARS-CoV were reported to have identical 3-D structures in the receptor-binding domain. SARS-CoV spike protein has a strong binding affinity to human Angiotensin-converting enzyme 2 (ACE-2) receptor, based on biochemical interaction studies and crystal structure analysis. SARS-CoV-2 and SARS-CoV spike proteins have high degree of homology and they share more than 70% identity in amino acid sequences^17^. Further, Wan et al., reported that glutamine residue at position 394 (E394) in the SARS-CoV-2 receptor-binding domain (RBD), corresponding to E479 in SARS-CoV, which is recognized by the critical lysine residue at pos-31 (K31) on the human ACE-2 receptor. Further analysis suggests that SARS-CoV-2 recognizes human ACE-2 receptor more efficiently than SARS-CoV increasing the ability of SARS-CoV-2 to transmit from person to person^18^. Thus, the SARS-CoV-2 spike protein was predicted to also have a strong binding affinity to human ACE-2 receptor.

ACE-2 is demonstrated as a functional SARS-CoV-2 spike (S) protein receptor *in-vitro* and *in-vivo*. It is required for host cell entry and subsequent viral replication. Zhou *et al.*,^19^ demonstrated that overexpressing ACE-2 receptor from different species in HeLa cells with human ACE-2, pig ACE-2, and civet ACE-2 receptor allowed SARS-CoV-2 infection and replication, thereby establishing that SARS-CoV-2 uses ACE-2 as a cellular entry receptor. In transgenic mice model with overexpression of human ACE-2 receptor, SARS-CoV infection enhanced disease severity and lung injury, demonstrating that viral entry into cells through ACE-2 receptor is a critical step^19, 20^. Thus for SARS-CoV-2 pathogenesis, spike (S) protein play a critical role by mediating entry of virus into the cell through human ACE-2 receptor and is an important target for vaccine development.

Here we report, design, production and pre-clinical testing of our DNA vaccine candidate. The proposed Coronavirus vaccine candidate comprises of a DNA plasmid Vector carrying spike (S) gene region of SARS-CoV-2 spike (S) protein along with gene coding for IgE signal peptide. The spike gene region was selected from submitted Wuhan Hu-1 isolate sequence (Genebank Accession No. MN908947.3). It’s expected that the plasmid construct with desired gene of interest will enter host cells, where it remains in the nucleus as an episome; without getting integrated into the host cell DNA. Thus using the host cell’s protein translation machinery, the inserted cloned DNA in the episome will direct the synthesis of the antigen it encodes. The approach involving the synthesis of antigen within the cells has several potential advantages. The protein produced by plasmid-transfected cells is likely to be expressed within the cell and folded in its native conformation. Further the signal peptide will prompt cells to translocate the protein, usually to the cellular membrane. The antigen is recognized by antigen presenting cells (APCs) and further induce antibodies including neutralizing antibodies through major histocompatibility complex (MHC) class pathway^13^. The DNA vaccine candidate induces antibody response against SARS-CoV-2 spike (S) protein, following immunization with just a single dose. Neutralizing antibody response was also demonstrated against wild type SARS-CoV-2 strain, which may play a substantial role in viral clearance and mitigation of human clinical disease. Immunogenicity of this DNA vaccine candidate targeting the SARS-CoV-2 S protein in animal model supports further clinical development of this candidate in response to the current COVID-19 pandemic situation.

## Material and Methods

### Selection of Spike (S) Gene region based on in-silico analysis

For our DNA vaccine candidate, the target antigen amino acid sequence of SARS-CoV-2 spike(S) from Wuhan Hu-1 isolate (Genebank Accession No. MN908947.3) was analysed *In-silico* by National Centre for Biotechnology Information (NCBI) blast tool and Clustal W multiple sequence alignment software to predict homology to other circulating SARS-CoV-2 spike(S) protein. In order to develop a vaccine candidate which provides broad protection against all circulating strains of SARS-CoV-2, antigenic region which is conserved and induce robust immune response is selected.

### Generation of ZyCoV-D vaccine construct

Gene sequence was submitted to GeneArt, Thermo Fisher Scientific and codon optimized full length Spike (S) region of SARS-CoV-2 virus with IgE signal sequence was synthesized. The chemically synthesized Spike(S) gene region preceded by IgE signal sequence was inserted into pVAX-1 plasmid DNA vaccine vector. Subsequently, the plasmid DNA construct was transformed in DH5-α™ chemically competent cells. After heat shock transformation step, *E. coli* clones carrying the plasmid DNA constructs were isolated by plating on LB agar plate containing Kanamycin antibiotic.

Single colonies were picked and inoculated in flasks containing LB broth from Hi-Media with Kanamycin. Flasks were incubated in 37°C incubator shaker at 225 rpm for 20 Hrs. Culture from each clone was used for plasmid isolation using miniprep plasmid isolation kit. Restriction digestion was carried out with *BamH1, Nhe1* and *Apa1* for all constructs to check expected band releases of inserts to select the positive clones. Positive clones were selected for preparation of glycerol stocks and stored at −70°C.

### *In-vitro* expression analysis of the constructs

In-vitro expression of DNA vaccine candidate was checked by transfection of the same in vero cell line. For transfection experiments, vero cells were seeded at density of 3× 10^5^ cells/ml in 6 well plates and kept in CO_2_ incubator to attain 80-90% confluency. After 24Hrs, once the cells reached the desired confluency, transfection was carried out in OptiMEM serum free medium with Lipofectamine 2000 reagent (Thermo Fisher). Two different concentrations (4μg and 8μg) of DNA construct was used for transfection experiments. After transfection, media was replenished with fresh DMEM media (Biowest) containing FBS. After 72Hrs, plates were fixed with 1:1 acetone and methanol Anti-S1 rabbit polyclonal antibody (Novus) was added to each well and incubated for 1Hr followed by incubation with FITC labelled anti-rabbit antibody (Merck). Fluorescence images were captured using an inverted microscope (ZeissAX10).

### Animal Immunization

The immunogenicity study for the ZyCoV-D vaccine was carried out in inbred BALB/C mouse, guinea pig, and New Zealand white rabbit model after having ethical approval from Institutional Animal Ethics Committee, CPCSEA Reg. No.: 335/PO/RcBi/S/01/CPCSEA, with IAEC approved application numbers: VAC/010/2020 and VAC/013/2020. BALB/c mouse (five to seven-week-old), guinea pigs (five to seven-week-old) and New Zealand White rabbits (six to twelve-week-old) were used in this study. For mouse intradermal immunization, on day 0; 25 and 100 μg of DNA vaccine was administered to the skin by using 31 gauge needle. Animals injected with empty plasmid served as vehicle control. Two weeks after immunization, animals were given first booster dose. Similarly all mice were given second booster dose two weeks after first booster dose. For guinea pig study, intradermal immunization was carried out using same dosing and schedule. In rabbits, DNA vaccine was administered to the skin by using needle free injection system (NFIS) at 500 μg dose at same 3 dose regimen and schedule. Blood was collected from animals on day 0 (before immunization) & 28 (after 2 dose) and on day 42 (after 3 dose) for immunological assessments from sera samples. In mouse model long term immunogenicity of the vaccine was assessed for up to day 126. Further, IFN-γ response from splenocytes at day 0, 28, and 42 were assessed.

### Measurement of antibody titres by ELISA

ELISA was performed to determine antibody titres in different animal sera samples. In brief, Maxisorp ELISA plates (Nunc) were coated with 50ng/well of recombinant S1 spike protein of SARS-CoV-2 (Acro, USA) in phosphate-buffered saline (PBS) overnight at 4 °C. Plates were washed three times with PBS then blocked with 5% skimmed milk (BD Difco) in PBS for 1 Hr at 37 °C. After blocking plates were then washed thrice with PBS and incubated with serial dilutions of mouse, guinea pig and rabbit sera and incubated for 2 Hrs at 37 °C. After that, plates were again washed thrice followed by incubation with 1:5,000 dilution of horse radish peroxidase (HRP) conjugated anti-guinea pig IgG secondary antibody (Sigma-Aldrich) or 1:2,000 dilution of HRP conjugated anti-mouse IgG secondary antibody (Sigma-Aldrich) or 1:5,000 dilution of HRP conjugated anti rabbit IgG secondary antibody (Sigma-Aldrich) for 1 Hr at 37 °C. After that again plates were washed thrice with PBS and then developed using TMB Peroxidase Substrate (KPL).Reaction was stopped with Stop Solution (1 N H_2_ SO_4_). Plates were read at 450 nm wavelength within 30 min using a multimode reader (Molecular Devices, USA).

### Virus neutralization assays using wildtype SARS-CoV-2

Micro-neutralization test (MNT) was performed at Translational Health Science and Technology Institute (THSTI), NCR Biotech Science Cluster, Faridabad – 121001 Haryana, India. The virus was obtained from the BEI resources, USA (Isolate USA-WA1/2020), passaged and titrated in Vero-E6 cells. The sera samples collected from immunized animals were heat-inactivated at 56 °C for 30 min followed by two fold serial dilution with cell culture medium. The diluted sera samples were mixed with a virus suspension of 100 TCID50 in 96-well plates at a ratio of 1:1 followed by 1 Hr incubation. This is followed by 1 Hr adsorption on Vero-E6 cells seeded 24 Hrs prior to experiment in 96 well tissue culture plate (1 × 10^4^ cells/well in 150 μl of DMEM +10% FBS). The cells were subsequently washed with 150 μl of serum free media and 150 μl of DMEM media supplemented with 2% FBS, followed by incubation for 3–5 days at 37°C in a 5% CO_2_ incubator. Cytopathic effect (CPE) was recorded under microscopes in each well. Neutralization was defined as absence of CPE compared to virus controls.

### Detection of neutralizing antibodies by competitive inhibition ELISA

Competitive inhibition ELISA was performed using SARS-CoV-2 neutralization antibody detection kit (Genscript). The kit detects circulating neutralizing antibodies against SARS-CoV-2 that block the interaction between the receptor binding domains of the viral spike glycoprotein (RBD) with the ACE2 cell surface receptor.

Different animal sera samples serially diluted with dilution buffer provided in the kit. The diluted sera samples were incubated with HRP conjugated RBD at 1:1 ratio for 30 min at 37°C along with positive and negative controls. The sera and HRP conjugated RBD mix was then added to the ELISA plate pre-coated with the ACE2 protein. After that, plates were incubated for 15 min at 37°C followed by washing four times with wash solution provided in the kit. After washing steps, TMB solution was added to the well and incubated in dark for 15 min at room temperature, followed by addition of stop solution. Plates were read at 450 nm. Inhibition concentration (IC_50_) of sera sample was calculated by plotting the percentage competition value obtained for each dilution verses serum dilution in a non-linear regression curve fit using Graph pad Prism 8.0.1 software.

### IFN-γ ELISPOT assay

For IFN-γ ELISPOT assay, spleens from immunized mice were collected in sterile tubes containing RPMI 1640 (Thermoscientific) media supplemented with 2X Antibiotic (Antibiotic Antimycotic, Thermoscientific). Cell suspensions were prepared by crushing the spleen with disk bottom of the plunger of 10 ml syringe (BD) in sterile petri plates. Then 5– 10ml of RPMI-1640 medium supplemented with 1X Antibiotic was added to it and the contents were mixed for homogeneity. Dishes were kept undisturbed for 2 min and the clear supernatant was pipetted out slowly into cell strainer (BD). The filtrate was collected in sterile tubes and the cells were pelleted by centrifugation at 4°C for 10min at 250×g in a centrifuge (Thermo Scientific). The pellet containing red blood cells (RBCs) and splenocytes were collected. 2-3 ml RBC Lysing Buffer (Invitrogen) was added to the pellet containing splenocytes and incubated at room temperature for 5-7 min. After incubation RPMI 1640 supplemented with 10% FBS (Biowest) and 1X antibiotic was added thrice the volume of RBC Lysing Buffer added previously. The pellets were washed with RPMI 1640 supplemented with 10% FBS and 1X antibiotic twice and were re-suspended in RPMI 1640 medium containing 10% FBS and 1X antibiotic and adjusted to a density of 2.0×10^6^ cells/ml. The 96 well Mouse IFN-γ ELISPOT kit (CTL, USA) plates pre-coated with purified anti-mouse IFN-γ capture antibody were taken out blocked with RPMI + 10% FBS +1X antibiotic for 1 Hr in CO_2_ incubator. The plate were then washed with PBS once and then 200,000 splenocytes were added to each well and stimulated for 24 Hrs at 37°C in 5% CO_2_ with pool of 12-mer peptides (GenScript) at a concentration of 5.0 μg/well spanning the entire SARS-CoV-2 S protein along with Negative control (RPMI 1640 supplemented with 10% FBS and 1X antibiotic and positive control (Concanavalin A, 1 μg /well). After stimulation, the plates were washed with PBS followed PBS containing 0.05% tween and spots were developed as per the manufacturer’s instructions provided along the kit. The plates were dried and the spots were counted on ELISPOT Reader S6 Versa, (CTL USA) and analysed with Immunospot software version 7.0.

### Biodistribution Study

Biodistribution of DNA vaccine candidate was studied in Wistar rat model. Two groups of Wistar rats received either a single bilateral, intradermal administration of 1 mg of plasmid or a single bilateral, intradermal administration of 0.5mg of plasmid per animal in each group respectively. At different time points (2 Hrs, 24 Hrs, 168 Hrs, and 336 Hrs) animals from each group were sacrificed and brain, lungs, intestine, kidney, heart, spleen, skin, and blood samples were harvested from the animals. All animal procedures were approved by the institutional animal ethics committee.

Total DNA was extracted from Wistar rat tissues samples using the DNeasy Blood & Tissue Kit (Qiagen, Germany) as per manufacturer’s instructions. Quantitative PCR was carried out with the PowerUp™ SYBR™ Green Master Mix (Applied Biosystems by Thermo Fisher Scientific, Lithuana) using plasmid specific primers on StepOne™ Real-Time PCR System (Applied Biosystems by Thermo Fisher Scientific, Lithuana). Plasmid DNA at different concentrations (1 x 10^2^ to 1 x 10^7^ copies/mL) were used to construct the standard curves The concentrations of samples were calculated by using standard curve with Applied Biosystems StepOnePlus™ software Ver. 2.2.2 (Applied Biosystems by Thermo Fisher Scientific, Lithuana).

### Statistical analysis

Statistical analysis of the results and graph creation were done with the Graph Pad Prism (version 8.0.1) and Microsoft Excel (version 7.0) for general statistical calculations, such as arithmetic mean and standard deviation. p values of <0.05 were considered significant.

## Results

### *In-silico* analysis of SARS-CoV-2 Spike (S) Protein

*In-silico* analysis confirmed more than 99% homology of the spike protein amino acid sequence from Wuhan strain with other circulating strains world including India.

### Generation of DNA vaccine constructs

Synthesis of SARS-CoV-2 spike (S) protein gene containing IgE signal peptide gene region and further cloning into pVAX-1® vector results in generation of SARS-CoV-2 DNA vaccine construct. Restriction digestion with *BamH1* resulting in linearized DNA fragments of ~6.8kb and restriction digestion analysis with *NheI* and *ApaI* resulting in generation of fragments of ~2.89kb of vector and ~3.89kb of spike protein (S) gene was used to confirm the insertion of spike(S) into the vector as shown in the Figure 1A and 1B. Gene sequencing analysis further confirmed the insertion of appropriate sequence in the desired orientation.

**Figure 1A-.**
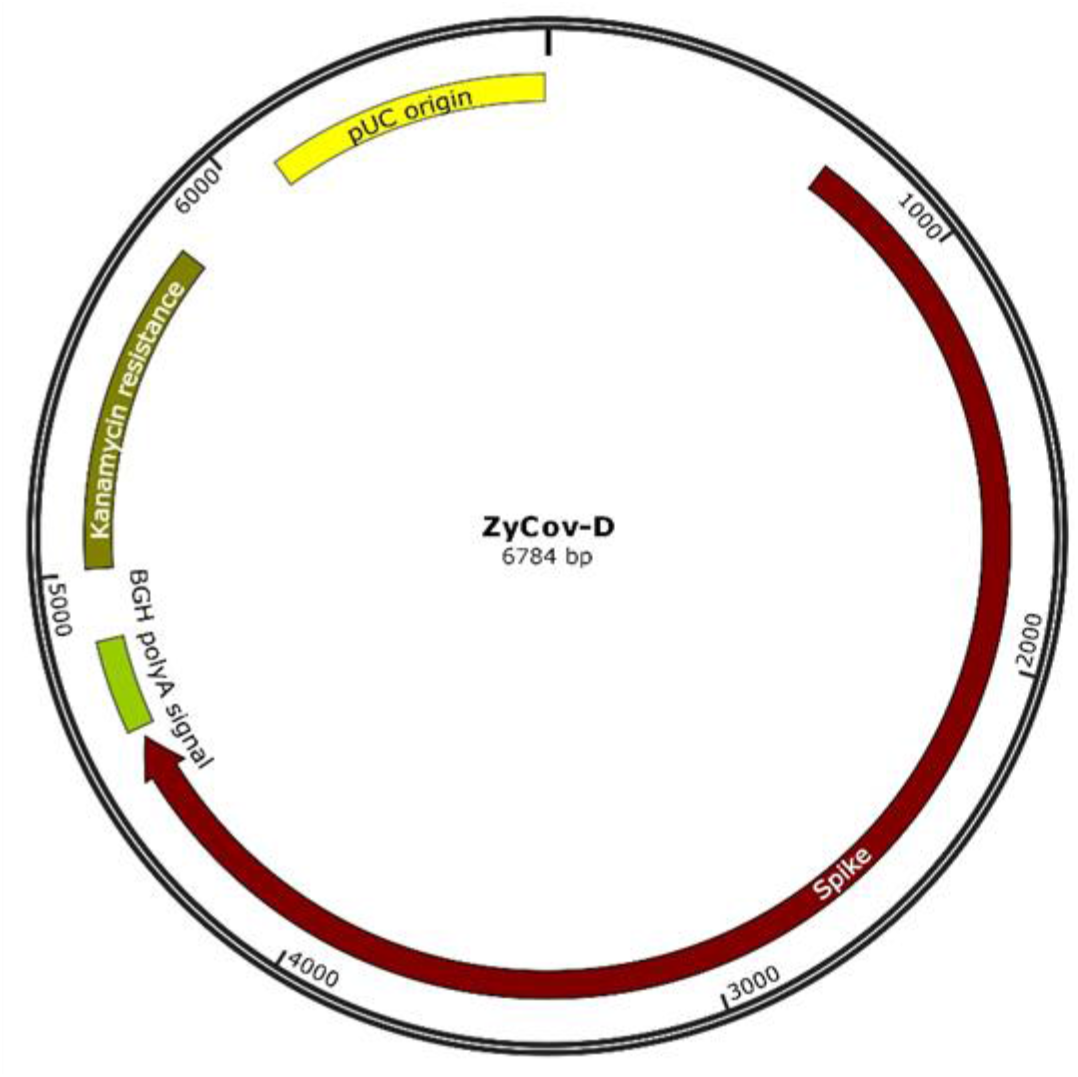
Schematic diagram of ZyCoV-D synthetic DNA vaccine constructs. pVAX-1vector containing SARS-CoV-2 spike gene insert

**Figure 1B-.**
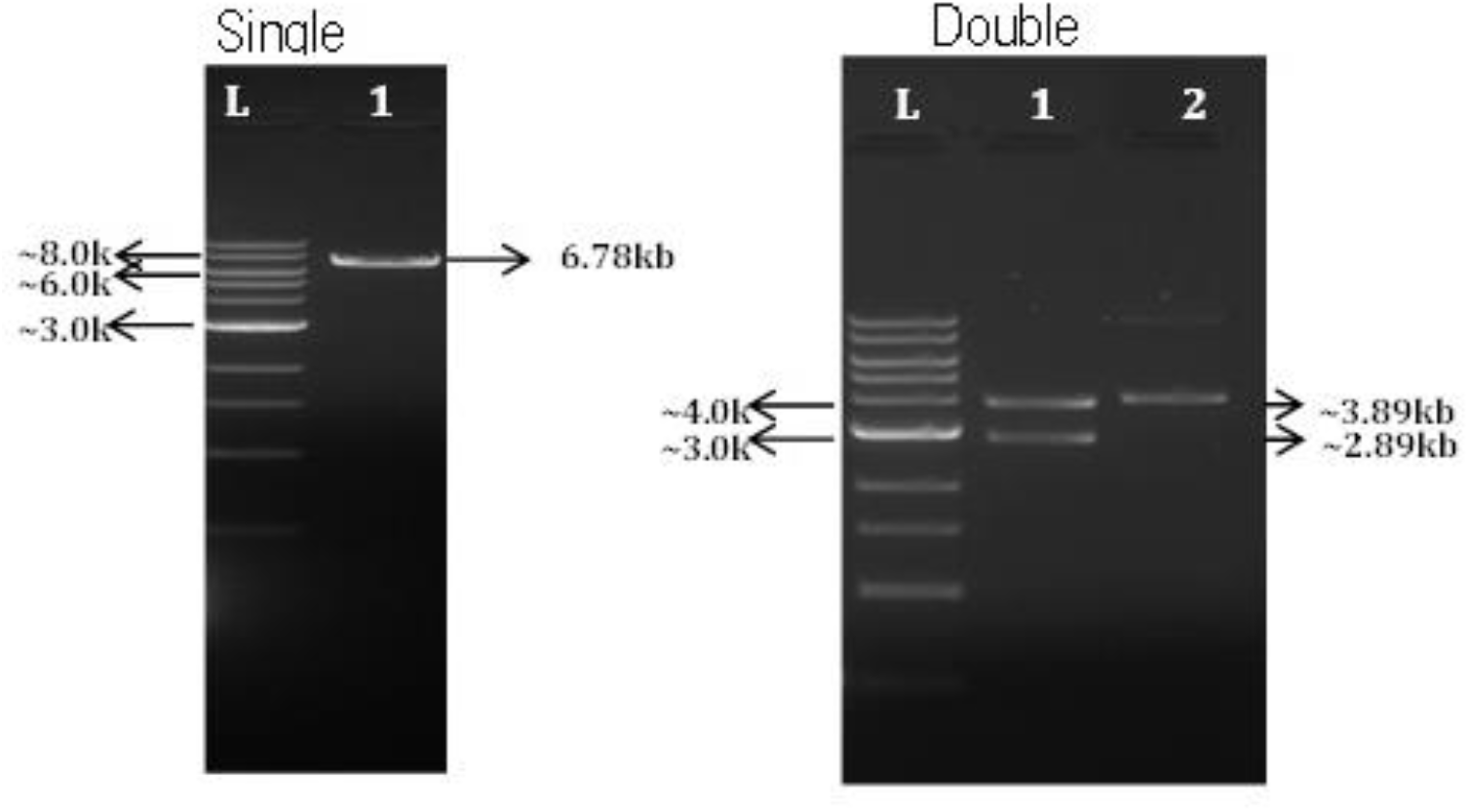
Restriction analysis of DNA vaccine constructs. Restriction digestion with *BamH1* resulting in generation of single fragments of ~6.78kb and restriction digestion of vector with *NheI* and *ApaI* resulting in generation of fragments of ~2.89kb of vector and ~3.89kb of spike protein (S) gene was used to confirm the insertion of spike(S) sequence into the vector

### In-vitro expression of DNA vaccine candidate

In Immunofluorescent studies were carried out to confirm the expression of S protein on candidate DNA vaccine transfected Vero cells (Figure 2). Immunostaining with FITC-labeled secondary antibody revealed the expression of the Spike protein after transfection of Vero cells with the candidate vaccine constructs.

**Figure 2-.**
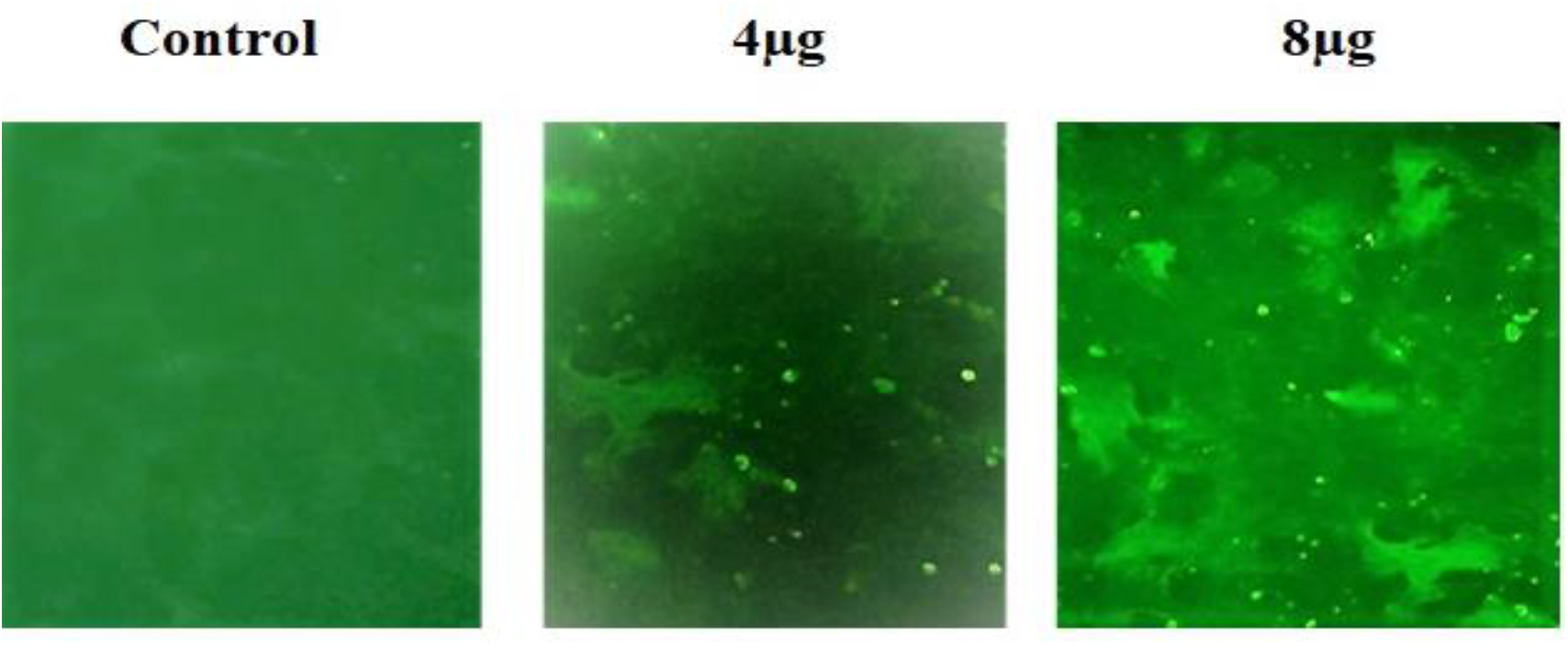
Analysis of *in-vitro* expression of Spike protein after transfection of Vero cells with DNA construct or empty plasmid (Control) by immunofluorescence. Expression of Spike protein was measured with polyclonal rabbit anti-SARS Spike Protein IgG and FITC anti-IgG secondary (green). Images were captured using inverted fluorescence microscope.

### Humoral immune response to DNA vaccine candidate

Immunization with DNA vaccine candidate by intradermal route elicited significant serum IgG responses against the S protein in doses-dependent manner in BALB/c mice, guinea pigs and rabbits with mean end point titres reaching ~28000 in BALB/C mice, ~140000 in guinea pigs and ~17000 in rabbits respectively on day 42 after 3 doses (Figure 3, 4 and Figure 5). Long term antibody response was studied in mice almost 3.5 months after the last dose and a mean end point IgG titres of ~18000 was detected (Figure 3) suggesting sustainable immune response was generated by DNA vaccine candidate.

**Figure 3-.**
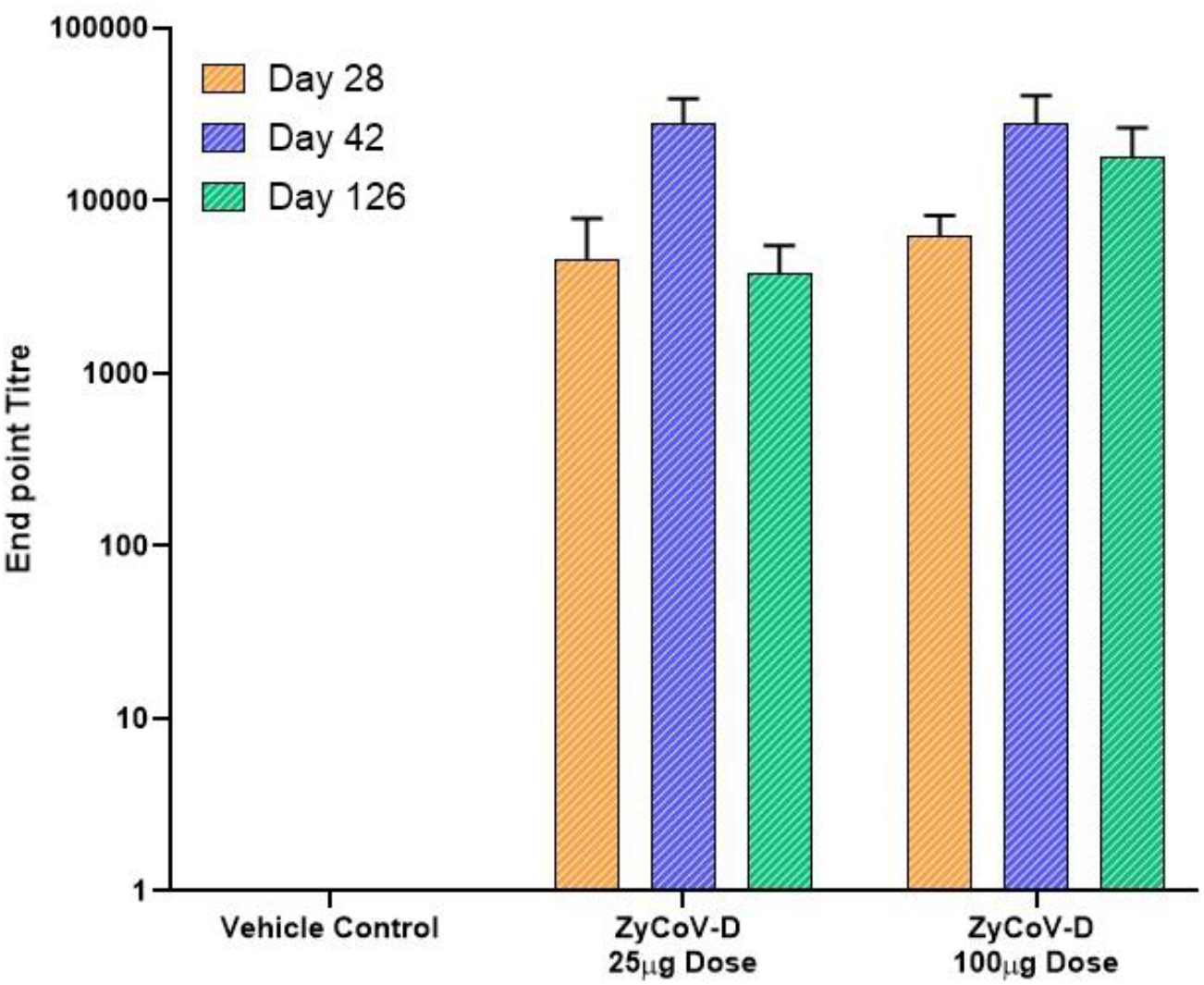
Antibody response after DNA vaccination in BALB/c mice and long term immunogenicity. BALB/c mice were immunized at week 0, 2 and 4 with DNA vaccine construct or empty control vector as described in the methods. Sera were collected at day 28 (orange), day 42 (blue) and day 126 (green) evaluated for SARS-CoV-2 S1-specific IgG antibodies.

**Figure 4-.**
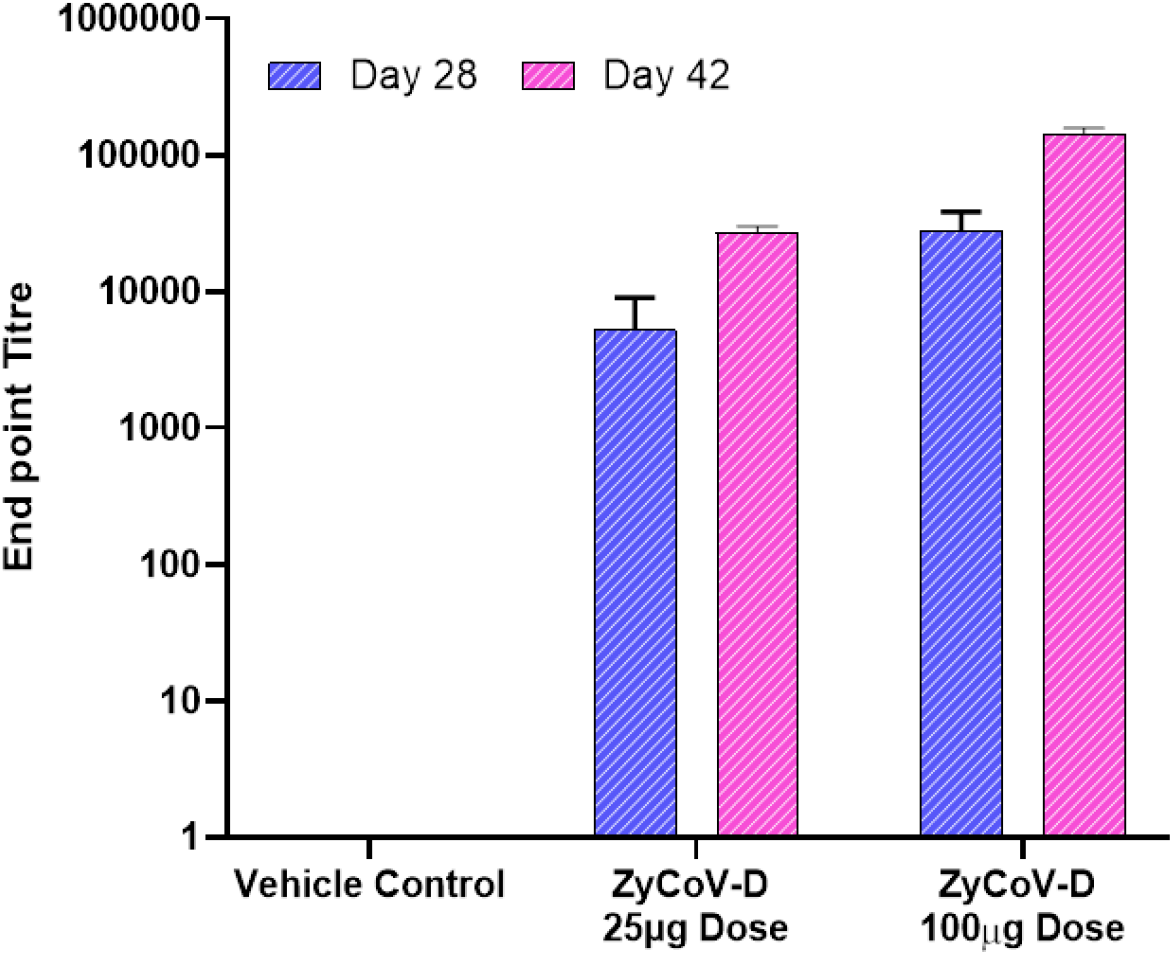
Antibody response after DNA vaccination in Guinea Pigs. Guinea pigs were immunized at week 0, 2 and 4 with DNA vaccine construct or empty control vector as described in the methods. Sera were collected at day 28 (blue) and day 42 (pink) and evaluated for SARS-CoV-2 S1-specific IgG antibodies.

**Figure 5-.**
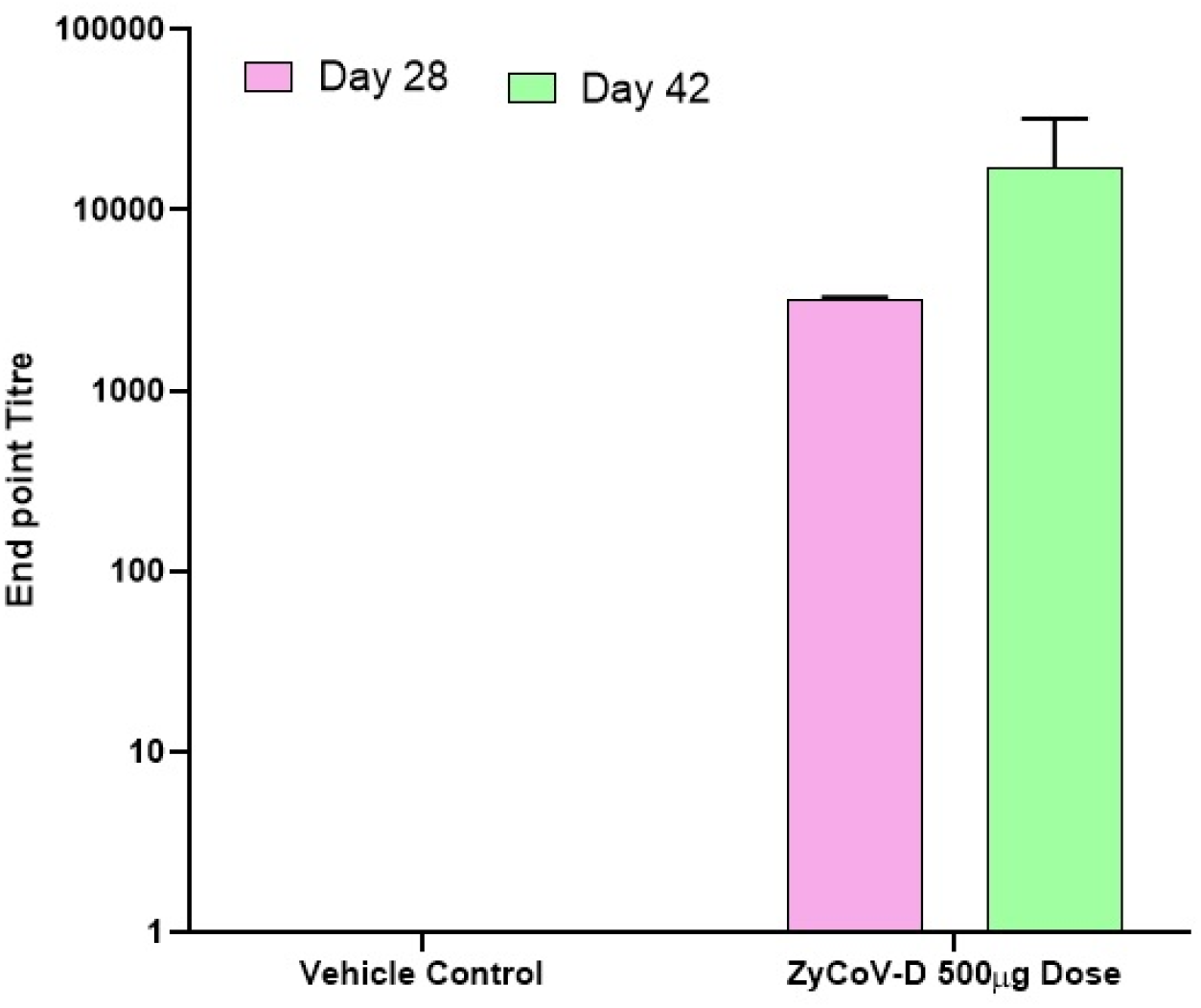
Antibody response after DNA vaccination in Rabbits. New Zealand White Rabbits were immunized at week 0, 2 and 4 with DNA vaccine construct or empty control vector as described in the methods. Sera were collected at day 28 (pink) and day 42 (green) and evaluated for SARS-CoV-2 S1-specific IgG antibodies.

Neutralizing antibody titres were evaluated in BALB/C mice, guinea pigs and rabbits following immunization by using micro-neutralization assay and Genscript neutralizing antibody detection kit. Neutralizing antibodies were elicited by DNA vaccine candidate in mice, guinea pigs and rabbits. Sera from DNA vaccine candidate immunized BALB/c mice could neutralize wild SARS-CoV-2 virus strains with average MNT titres of 40 and 160 at day 42 with 25 and 100 μg dose regimens respectively (Table 1). Using Genscript neutralizing antibody detection kit average IC_50_ titres of 82 and 168 were obtained at day 42 with 25 and 100 μg dose regimens respectively. Further, neutralizing antibodies were also detected in long term immunogenicity studies in BALB/c mice. Significant rise in neutralizing antibodies levels were also observed in guinea pigs and rabbits (Table 1).

**Table 1-.**
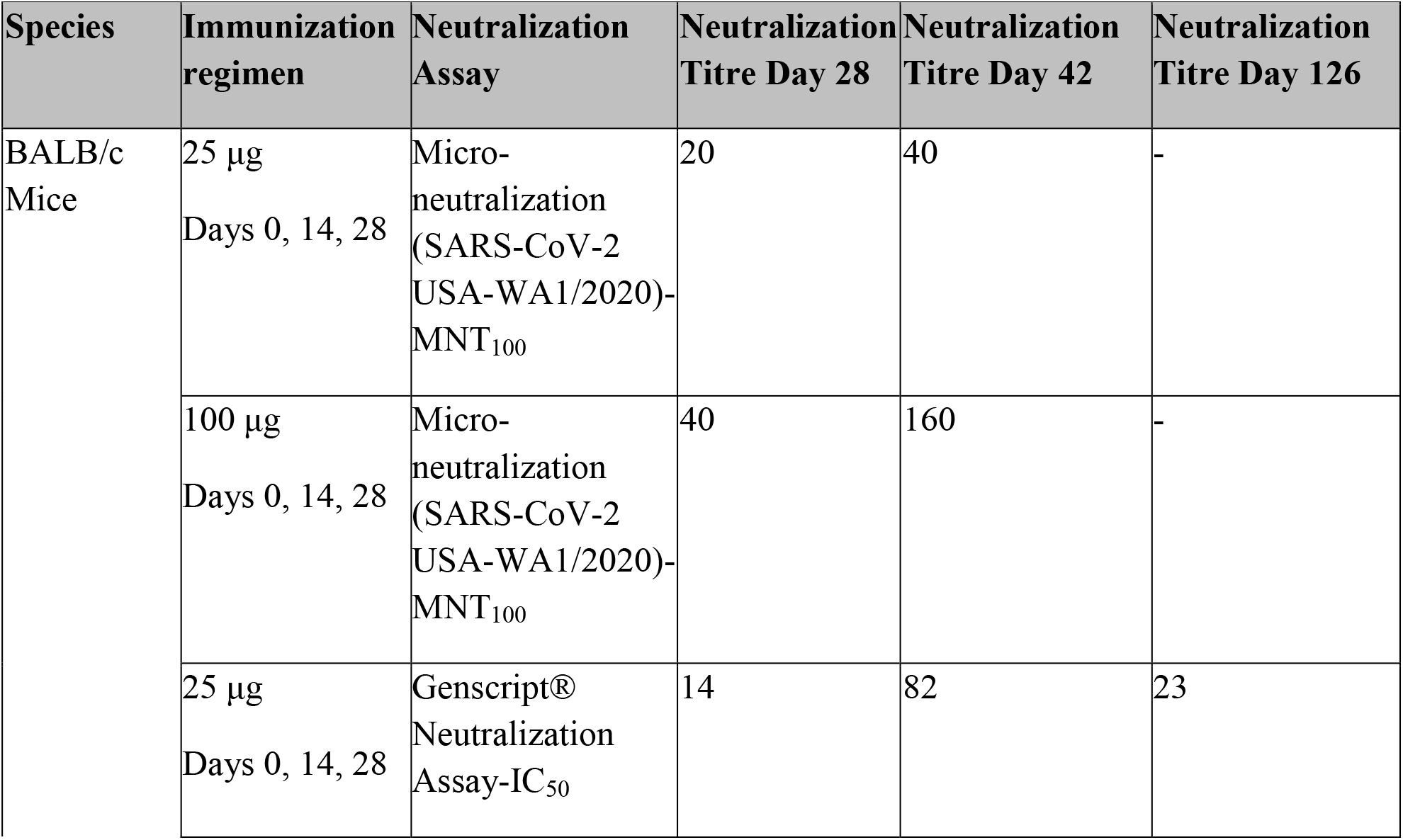
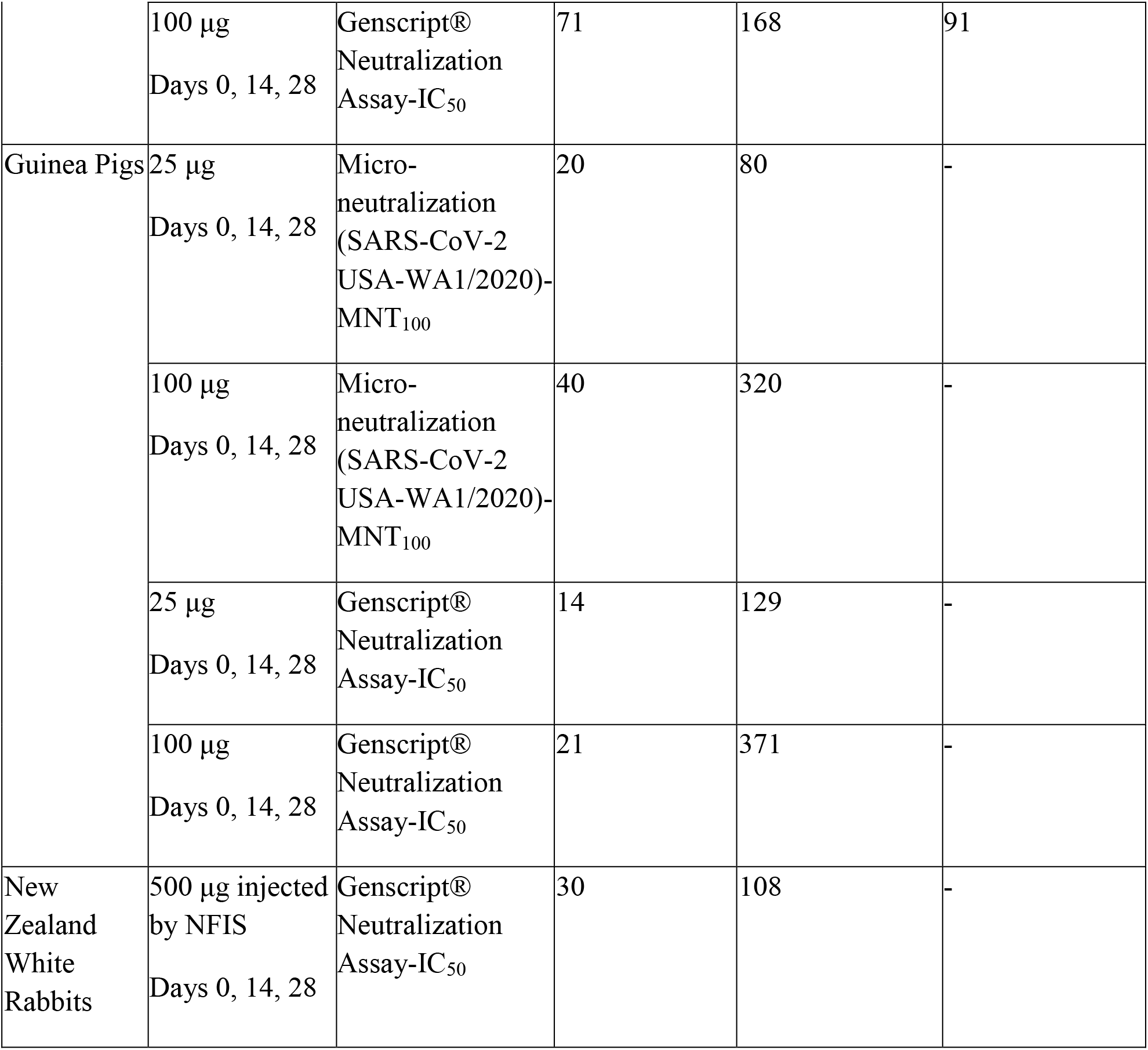
Sera neutralizing antibody titres after DNA vaccine administration to BALB/c mice, guinea pigs and rabbits.

### Cellular immune response to DNA vaccine candidate

T cell response against SARS-CoV-2 spike antigen was studied by IFN-γ ELISpot assay. Groups of BALB/c mice were sacrificed at day 14, 28, 42 post-DNA vaccine administration (25 and 100 μg dose). Splenocytes were harvested, and a single-cell suspension was stimulated for 24 h with pools of 12-mer overlapping peptides spanning the SARS-CoV-2 spike protein. Significant increase in IFN-γ expression, indicative of a strong Th1 response, of 200-300 SFC per 10^6^ splenocytes against SARS-CoV-2 spike peptide pool was observed for both the 25 and 100μg dose in post 42 day immunized mice splenocytes (Figure 6).

**Figure 6-.**
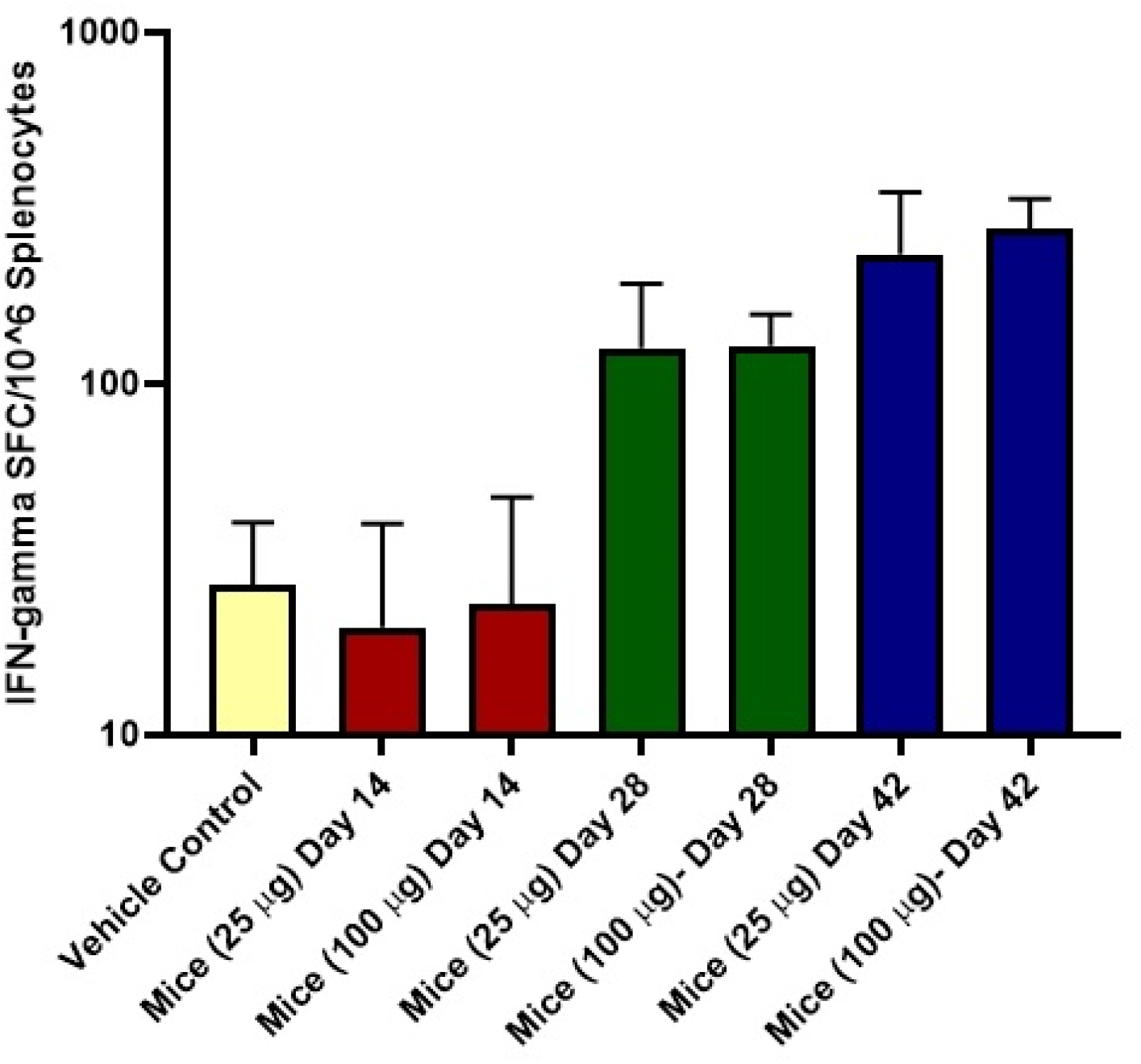
Detection of IFN-γ responses in BALB/c mice post-administration of DNA vaccine. BALB/c mice were immunized with 25 and 100 μg of DNA vaccine. IFN-γ responses were analyzed in the animals on days 14, 28 and 42. T cell responses were measured by IFN-γ ELISpot in splenocytes stimulated for 24h with overlapping peptide pools spanning the SARS-CoV-2 spike region

### Bio distribution

Biodistribution of DNA vaccine candidate was evaluated in Wistar Rats at single dose of 0.5mg and 1.0 mg administered intradermaly. DNA was extracted from various tissue samples including the site of injection, brain, blood, lungs, intestine, kidney, heart and spleen at different time points post injection as described above. RT-PCR was performed by plasmid specific primers to detect copy numbers. Following injection of maximum ~ 10^14^ plasmid DNA copies in Wistar Rats, maximum local concentration of 10^3^ –10^7^ plasmid copies at the site of injection were detected two hours post injection. (Figures 7A and 7B). We also observed bio-distribution of plasmid molecule in blood, lungs, intestine, kidney, heart, spleen, skin post 24 Hrs of injection which got cleared by 336 Hrs (day 14) in most of the organs except skin (site of injection) where only 10^2^-10^3^ copies were detected. Although by 672 Hrs (day 28) post injection, no plasmid copies were detected at site of injection as well.

**Figure 7A-.**
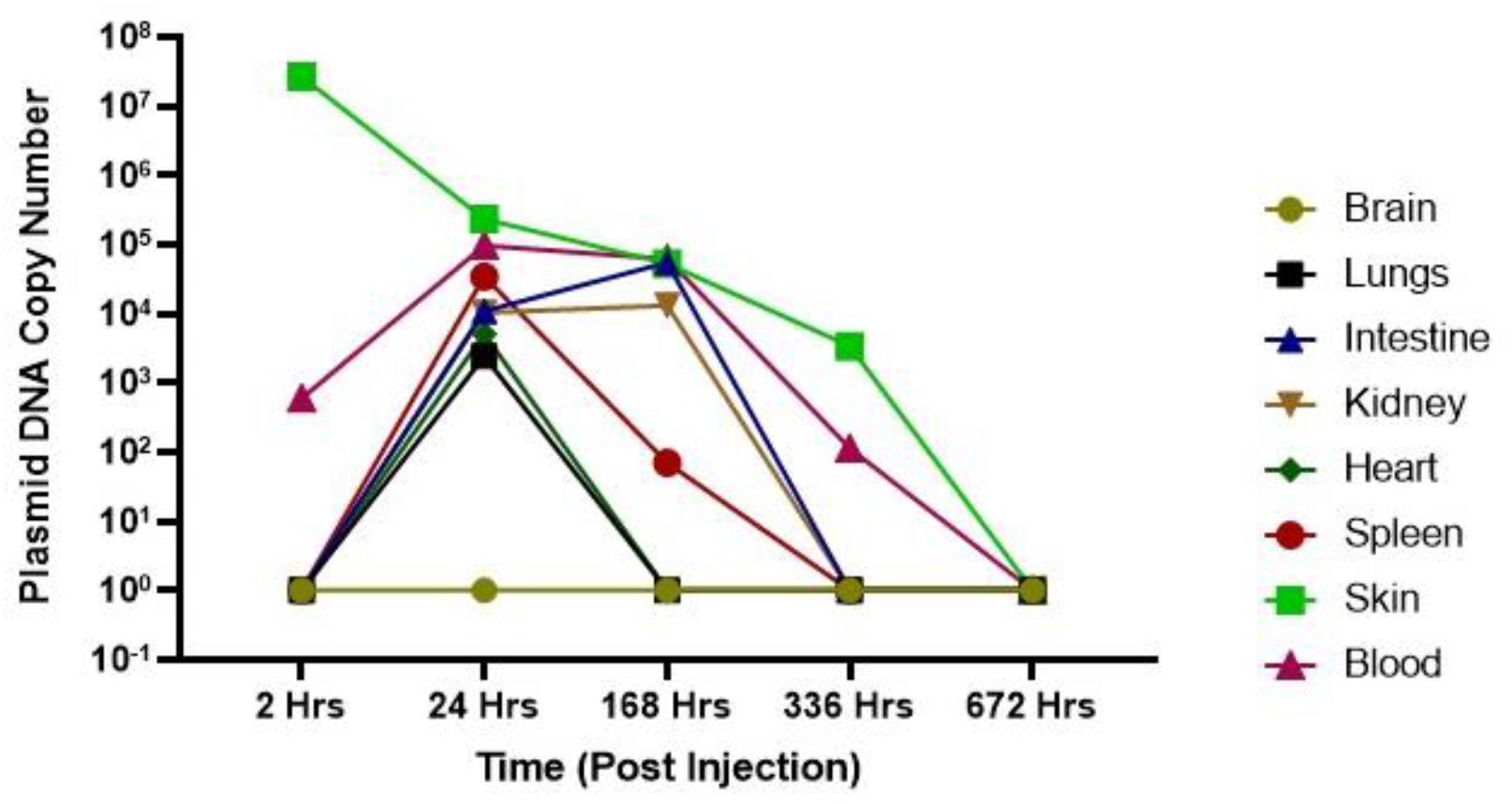
Bio-distribution of DNA vaccine after 1mg dose. Plasmid copy levels as determined by RT-PCR, in the tissue samples at site of injection, brain, blood, lungs, intestine, kidney, heart and spleen at different time point after injection in animals.

**Figure 7B-.**
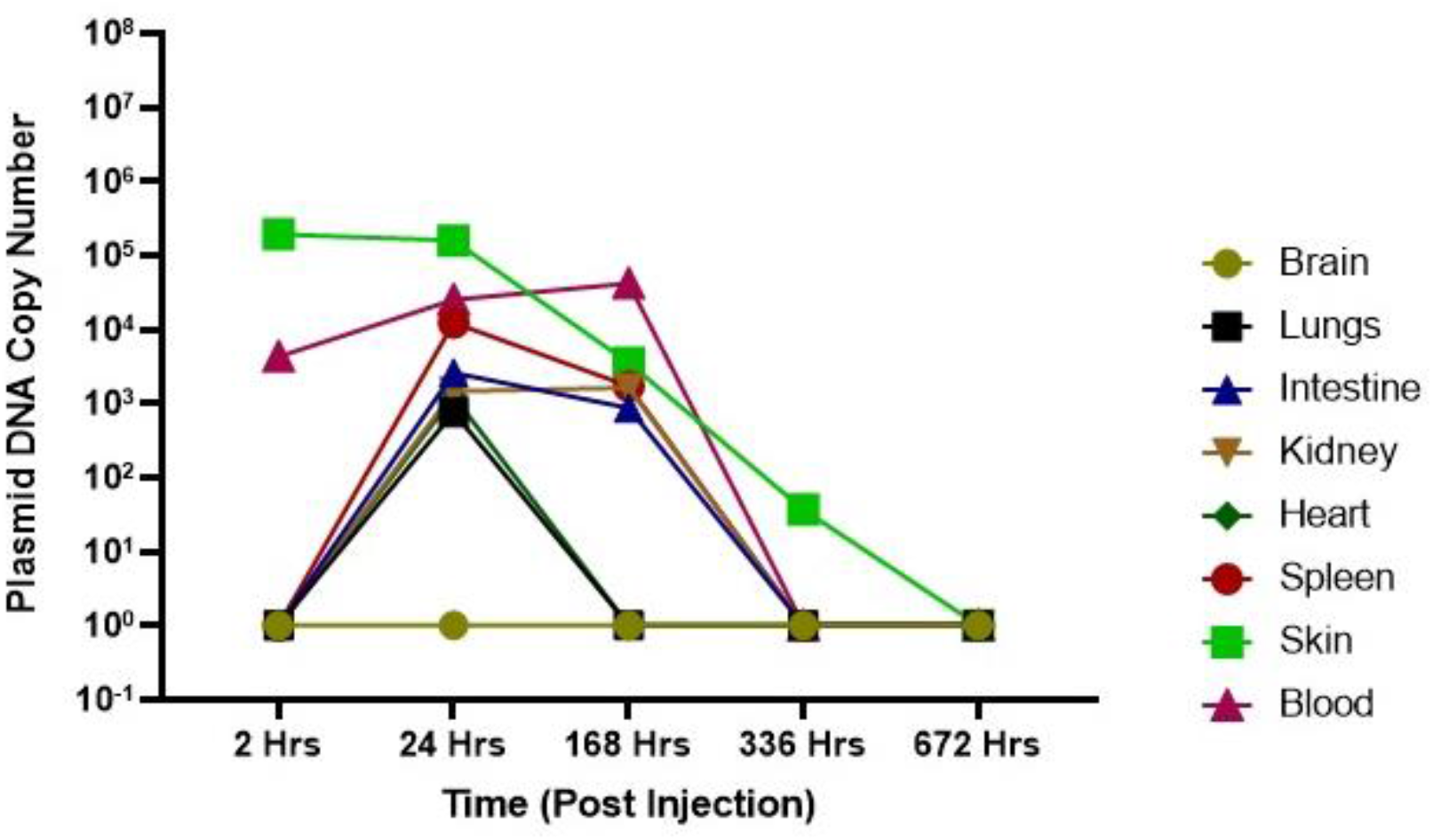
Bio-distribution of DNA vaccine after 0.5mg dose. Plasmid copy levels as determined by RT-PCR, in the tissue samples at site of injection, brain, blood, lungs, intestine, kidney, heart and spleen at different time point after injection in animals.

## Discussion

Development of safe and effective vaccine against SARS-CoV-2 is need of hour to curb the global pandemic. DNA vaccine platform has several advantages, which positions it well to respond to disease outbreaks, such as COVID-19. The ability to design and immediately synthesize candidate vaccine constructs allow us to carry out *in-vitro* and *in-vivo* testing within days of receiving the viral sequence. The expression and localization of S protein expressed by ZyCoV-D were investigated using an immunofluorescence assay. The immunofluorescence assay with rabbit anti–S1 antibody revealed a strong signal in the vero cells transfected with ZyCoV-D. In contrast, the positive signal was not detected in cells transfected with control vector. This demonstrates the ability of the ZyCoV-D vaccine to express strongly in mammalian cells and that antibodies induced by this construct can bind their target antigen. Further, the DNA plasmid manufacturing process is easily scalable with substantial yields, and has the potential to overcome the challenges of conventional vaccine production in eggs or cell culture. Additionally, we have also studied stability profile of our vaccine candidate (unpublished data). The stability data suggests that our DNA vaccine candidate can be stored at 2–8 °C for long term and further at 25°C for short term (few months). In the context of a pandemic outbreak, the stability profile of a vaccine plays a vital role easy deployment and distribution for mass vaccination. Further, we will like to highlight that ZyCoV-D was developed using a pVAX-1 vector, which has been used in number of other DNA vaccines in past and have been proven to be very safe for human use ^14,21^.

ZyCoV-D was evaluated *in-vivo* in different animal models and has demonstrated ability to elicit immunogenic response against SARS-CoV-2, S-antigen in animal species. Primary antibody response starts mounting in serum two weeks after two doses and reaches pick two weeks after third immunization. The serum IgG levels against spike antigen in mice were maintained even after three months post last dosing suggesting a long-term immune response generated by the DNA vaccine candidate. This also indicates that ZyCoV-D can possibly induce robust secondary anamnestic immune response upon re-exposure, generated by balanced memory B and helper T cells expression and has been reported for other DNA vaccine candidates^22^.

We reported serum neutralizing (Nab) titres following DNA vaccination, which was tested by micro-neutralization assay and Genscript neutralizing antibody detection kit. The Nab titre values tested by both methods demonstrated that the DNA vaccine candidate generates robust response and neutralizes the SARS CoV-2 virus conferring protective immunity against infection. In future if these Nab titres will be established as correlate of protection across multiple vaccine studies in both animals and humans, then this parameter can be utilized as a benchmark for clinical development of SARS-CoV-2 vaccines.

We also observed that ZyCoV-D vaccine is capable of inducing T-cell response complementary to antibody response in mice model as demonstrated by IFN-γ ELISPOT. This is very important as successful DNA vaccination is known to induce both humoral and cellular responses in both animals and human ^14,15,16,23^. The mechanism of action for DNA vaccine candidate includes both class-I antigen-processing pathways (i.e., intracellular processing of viral proteins and subsequent loading onto MHC class-I molecules) and class-II antigen-processing pathways (i.e., endosomal loading of peptides generated from endocytosed viral antigens secreted from cells using IgE signal peptide onto MHC class II molecules). Among T cell responses, a balanced Th1/Th2 response is important because vaccine-associated enhanced respiratory disease (VAERD) is associated with Th2-biased immune response. Indeed, immunopathologic complications characterized by Th2-biased immune responses have been reported in animal model of the SARS-CoV or MERS-CoV challenge ^24,25,26^ and similar phenomena have been reported in clinic trials vaccinated with whole-inactivated virus vaccines against RSV and measles virus ^27,28^. In addition, the importance of Th1 cell responses has been highlighted by recent study of asymptomatic and mild SARS-CoV-2 convalescent samples^29^. These results collectively suggest that vaccines capable of generating balanced antibody responses and Th1 cell responses may be important in providing protection against SARS-CoV-2 diseases.

The usefulness and efficiency of a spring-powered, needle-free Injection System (NFIS) for delivering ZyCoV-D vaccine in rabbits was also demonstrated in the study. Similar observation was reported earlier with application of NFIS for DNA vaccines against Hantavirus and Zika virus ^30,31^. The use of NFIS eliminates use of needles during vaccine administration thus eliminates the costs and risk associated with sharp-needle waste. Further, NFIS doesn’t required external energy sources such as gas cartridges or electricity and spring provides the power for the device. These injector create a stream of pressurized fluid that penetrates upto 2 mm in skin at high velocity resulting in uniform dispersion and higher uptake of DNA molecules in cells compare to needle and syringe where the intradermal accumulation is inconsistent across individuals (as measured by bleb size) and varies among animal species^30^.

Bio-distribution pattern for ZyCoV-D was also evaluated and level of plasmid DNA was measured at different intervals in various tissues in Wistar rats post intradermal injection. Post intradermal injection, the plasmid was found to clear off from most of the organs by 14 days post injection except site of injection, which also cleared off by 28 days post injection. Our outcome was very similar to other DNA vaccine candidate including HIV-1, Ebola, Severe Acute Respiratory Syndrome (SARS), and a West Nile Virus candidate developed^32^. Biodistribution and plasmid copy number detection studies for the vaccine candidates were done in different animal models^32^. It was observed that animals injected with 2mg (equivalent to 10^14^ plasmid copies) by both intramuscular and subcutaneous route have detectable plasmid copies in first one week of vaccination at the site of injection with copies in order of 10^4^-10^6^. Over the period of 2 months, the plasmid clears from the site of injection with only a small percentage of animals in group (generally 10-20%) retaining few copies (around 100 copies) at the injection site. Directly after injection into skin or muscle, low levels of plasmids are transported via the blood stream and detected in various organs at early time points. However, the plasmids are eventually, cleared from the organs and are normally found exclusively at the site of injection at later time points.

In summary, these initial results demonstrate the immunogenicity of our ZyCoV-D DNA vaccine candidate in multiple animal models. These studies strongly support the clinical evaluation as a vaccine candidate for COVID-19 infection.

## Acknowledgments

Authors acknowledge the support from PharmaJet, Inc. Golden, CO, USA for providing us PharmaJet® Tropis® Needle-Free Injection System (NFIS) for vaccine delivery.

## Funding

Development of ZyCoV-D was supported by a grant-in-aid from Covid-19 Consortium under National Biopharma Mission, Department of Biotechnology, Government of India, to Cadila Healthcare Ltd. (Grant no. BT/COVID0003/01/20).

## Authors Contribution

KM conceptualize, design, develop the vaccine candidate, provide guidance on data analysis and manuscript preparation, AD design, develop vaccine candidate, perform data analysis for ELISPOT assay and prepare manuscript, CRTM and HC develop analytical procedures for testing of the vaccine and perform data analysis for ELISA, neutralization and biodistribution assay, HPRP, HSC, GS, PD, SV, SP, SR developed process for vaccine production, MB perform animal experiment, AS perform ELISPOT assay, NL and MAR perform ELISA assay, SK, AS, VS and AP design and developed clone.

## Notes

### Competing Interest Statement

The authors have declared no competing interest.

